# Generation and characterization of Induced Pluripotent Stem Cell models with *SMAD3* mutations to study the molecular mechanism of Loeys-Dietz Syndrome Type III aneurysm formation

**DOI:** 10.1101/2025.08.26.667399

**Authors:** Nathalie P. Vroegindeweij-de Wagenaar, Janette van der Linden, Hanny J.H.M. Odijk, Matthijs Snelders, Ingrid M.B.H. van de Laar, Roland Kanaar, Ingrid van der Pluijm, Jeroen Essers

**Affiliations:** Department of Molecular Genetics, Oncode Institute, Erasmus MC, University Medical Center Rotterdam, Rotterdam, The Netherlands; Department of Cardiology, Erasmus MC, University Medical Center Rotterdam, Rotterdam, The Netherlands; Department of Farmacology, Erasmus MC, University Medical Center Rotterdam, Rotterdam, The Netherlands; Department of Clinical Genetics, Erasmus MC, University Medical Center Rotterdam, Rotterdam, The Netherlands; Department of Vascular Surgery, Cardiovascular Institute, Erasmus MC, University Medical Center Rotterdam, Rotterdam, The Netherlands; Department of Radiotherapy, Erasmus MC, University Medical Center Rotterdam, Rotterdam, The Netherlands

## Abstract

Loeys-Dietz Syndrome type 3 (LDS3) is caused by pathogenic (P)/likely pathogenic (LP) variants in the *SMAD3* gene and is characterized by aneurysm formation and arterial tortuosity, which can lead to life-threatening complications. There is an unmet need for suitable cell models to study LDS3 at a cellular and molecular level. Induced pluripotent stem (iPS) cells offer a promising approach because they can be genetically modified using CRISPR/Cas9 technology and differentiated into disease-relevant cell types. As it is difficult to obtain aortic vascular smooth muscle cells (VSMCs) from patients, iPS cells differentiated into VSMCs provide an ideal model to study cellular aneurysmal phenotypes.

In this study, we generated iPS cell models carrying (P/LP) *SMAD3* variants. These cell models were generated either by using CRISPR/Cas9 mediated introduction of indels and deletions to introduce *SMAD3* variants, or by reprogramming of fibroblasts derived from *SMAD3* patients. These iPS cell lines were characterized for SMAD3 expression by Western blotting and validated for pluripotency through immunofluorescence and qPCR. Moreover, the patient-derived iPS cell lines were shown to differentiate into smooth muscle cells (SMCs), which are relevant to study the molecular mechanisms underlying aneurysm formation in LDS3 patients. Our findings highlight the potential of these iPS-based models to investigate the pathophysiology of LDS3 and facilitate the development of therapeutic strategies for aortic aneurysms.

## Introduction

Autosomal dominant P/LP variants in the *SMAD3* gene have been implicated in Loeys-Dietz Syndrome type 3 (LDS3), also known as Aneurysm Osteoarthritis Syndrome (AOS) [1]. The most common cardiovascular features of LDS3 are aneurysms and vascular tortuosity of the aorta and middle sized arteries, but skeletal and joint abnormalities, including osteoarthritis, are also frequently observed [2, 1]. Dissection or rupture of aneurysms can result in severe morbidity and even mortality [3].

Over 60 different P/LP variants in *SMAD3* have been identified as causing for LDS3 [4-8]. The clinical phenotype of affected individuals is highly variable, and, at present, the only treatment option for aneurysms is surgical intervention and medication to reduce hemodynamic stress. To develop more targeted therapies, a better understanding of the underlying disease mechanism is essential. Critical to achieving this understanding is the development of reliable cellular models to study the disease at a cellular level. Vascular smooth muscle cells (VSMCs) are the primary cell type affected in aortic aneurysmal tissue. However, obtaining VSMCs from patients is challenging, as they can only be isolated from aortic aneurysm tissue which is available in limited amounts. Induced pluripotent stem (iPS) cells offer a promising alternative model, as they can be derived from patient fibroblasts or blood and can be differentiated into SMCs. The generation of iPS cells involves reprogramming of somatic cells to a pluripotent state by introducing genes such as *OCT4, SOX2, KLF4*, and *C-MYC*. This method is less invasive than obtaining tissue biopsies and iPS cells can be differentiated into disease-relevant cell types.

One challenge when studying specific P/LP variants in patient-derived material is the limited number of patients with specific variants, which makes it difficult to exclude variation due to external factors such as genetic background. To mitigate this, introducing mutations with CRISPR/Cas9 in iPS cells derived from healthy individuals can generate isogenic control lines with the mutation of interest.

This approach enables more accurate studies of the LDS3 phenotype by providing a mutation-specific model, as well as by introducing multiple *SMAD3* mutations within the same isogenic background, free from confounding factors.

In this paper, we generated iPS-based cell models for *SMAD3* mutations using CRISPR/Cas9-mediated gene editing and by reprogramming fibroblasts from LDS3 patients. We validated SMAD3 protein expression in these lines through Western blot analysis. Furthermore, we differentiated the patient-derived iPS cells into smooth muscle cells (SMCs), which are critical for studying the molecular mechanisms underlying LDS3. Our study highlights the potential of iPS-based models to study *SMAD3* variants and provides a foundation for further research into LDS3 disease mechanisms and potentially also therapeutic development.

## Materials and methods

### Cell culture

HEK293 cells and fibroblasts were cultured in Dulbecco’s Modified Eagle’s Medium (DMEM, Gibco, 11965-092) supplemented with 10% fetal calf serum (FCS, Capricorn, FBS-12A) and 1% penicillin/streptomycin (P/S, Sigma, P0781). iPS cells were cultured on Matrigel (354277, Corning™) coated dishes in mTeSR (plus) medium (Stemcell Technologies). iPS cells were passaged as aggregates with ReLeSR (Stemcell Technologies). iPS cells were dissociated as single cells using Accutase (Stemcell Technologies). All cells were incubated at 37°C with 5% CO_2_.

### sgRNA design, plasmids and primers

sgRNAs were designed with the Zhang Lab CRISPR Tool (Zhang Lab MIT, 2017) to target exon 6, either to create indels or to target both upstream and downstream sequences of the *SMAD3* gene to delete an entire *SMAD3* allele. Short overhangs were added to the sequence to facilitate cloning into the plasmids, and these sgRNAs were ordered from IDT Technologies (Supplementary Table 1). sgRNAs to create indels were cloned into plasmid eSpCas9(1.1)-T2A-EGFP (Addgene, 71814 with T2A-EGFP cloned in) and sgRNAs to delete the entire *SMAD3* allele were cloned into plasmid pSpCas9(BB)-2A-Puro (PX459) V2.0 (Addgene, 62988) according to a previously published protocol [9]. pEGFP-N2 (Clontech, 6081-1) was used as a transfection control. Primers were designed for the DNA sequence surrounding the target locations of the sgRNAs to be able to analyze targeting by PCR and sequencing. Primers were ordered at IDT Technologies (Supplementary Table 2, Figures 1 and 2).

**Figure 1.**
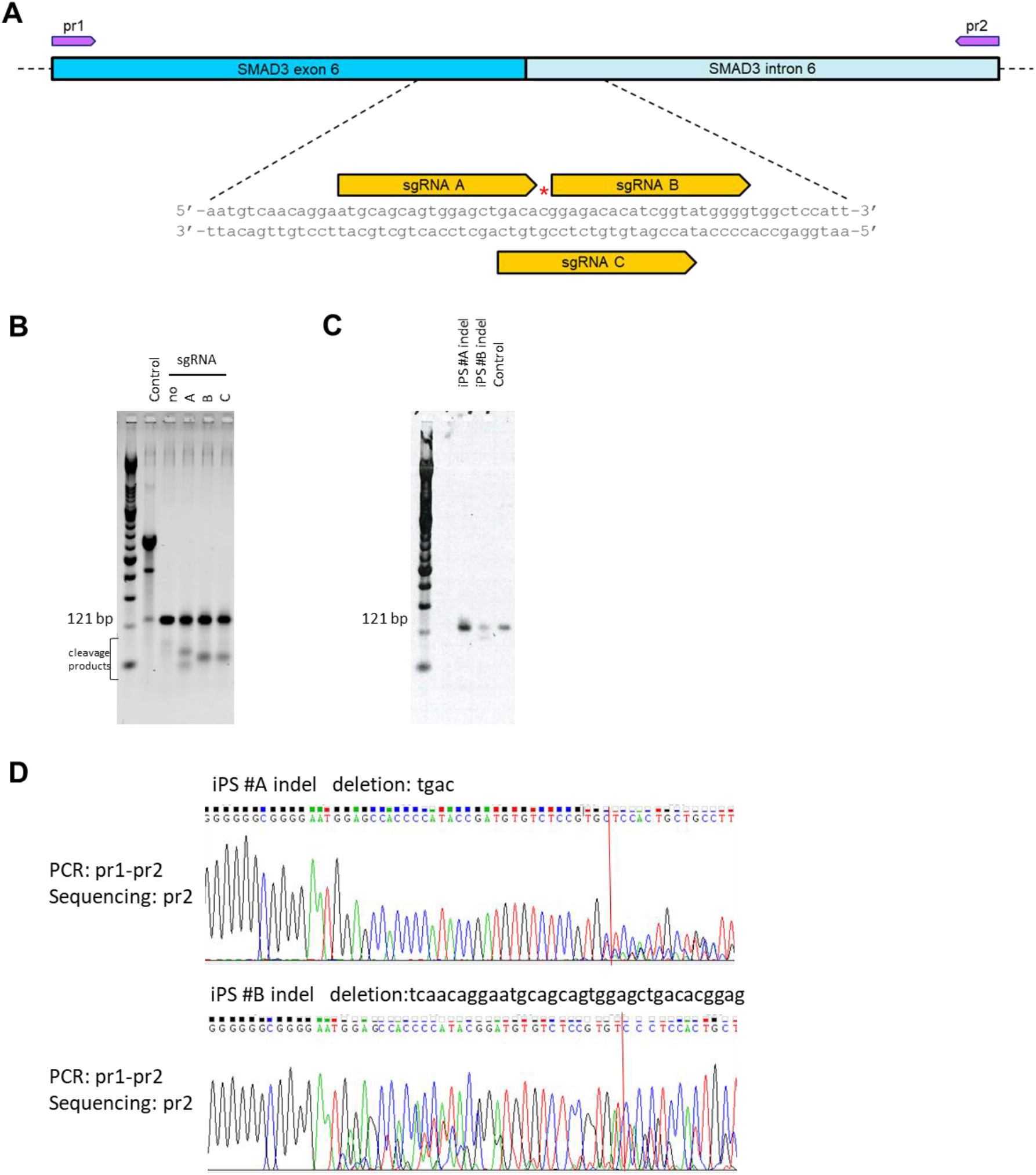
Generation and validation of *SMAD3* indel mutations using CRISPR/Cas9. (A) Strategy to introduce small deletions in *SMAD3* using CRISPR/Cas9. A schematic representation of exon 6 and the adjacent intronic region of the *SMAD3* gene is shown. Three different sgRNAs (sgRNA-A, sgRNA-B, sgRNA-C) were designed. The red star indicates the location of a known *SMAD3* mutation. Primer binding sites for PCR and sequencing (pr1 and pr2) are indicated. (B) Surveyor assay to assess sgRNA efficiency. Lanes show a 100 bp marker, a positive control, and the Surveyor assay performed on PCR products (pr1-pr2) from DNA of cells transfected with Cas9 alone (no sgRNA) or with sgRNA-A, sgRNA-B, or sgRNA-C. The presence of cleavage products, smaller than the PCR product, indicates the formation of heteroduplexes due to indels introduced by CRISPR/Cas9. (C) PCR genotyping of iPS cell clones. Lanes show a 100 bp marker, PCR amplification of the *SMAD3* locus (pr1-pr2) from genomic DNA of two mutant iPS cell clones and control iPS cells. (D) Sanger sequencing confirming CRISPR/Cas9-induced mutations in iPS clones. PCR amplification was performed using pr1-pr2, with sequencing carried out from the pr2 primer. The red line marks the Cas9 cut site. Clone iPS #A indel harbors a 4 bp deletion, while clone iPS #B indel contains a 34 bp deletion.

**Figure 2.**
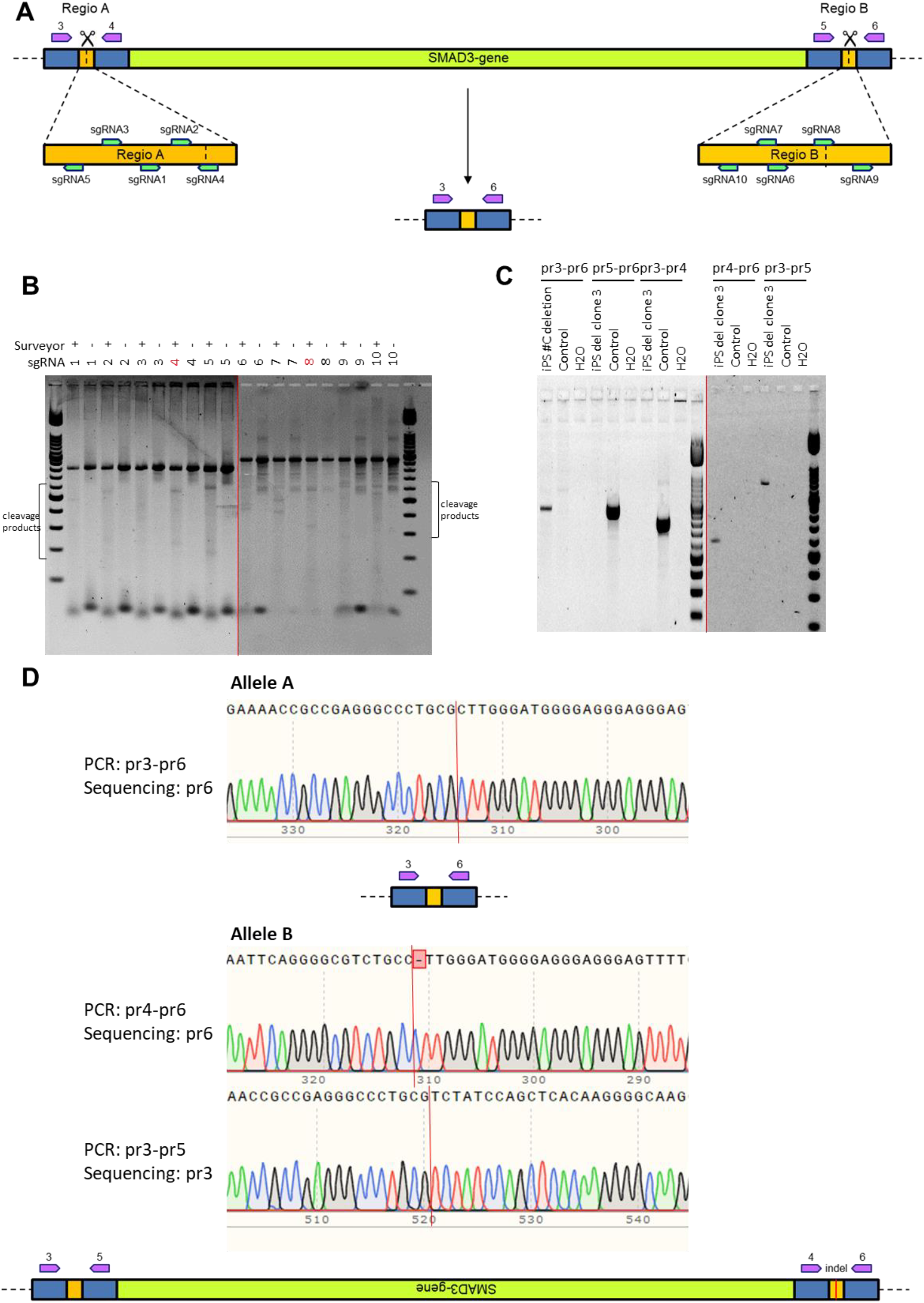
Generation of *SMAD3* knockout (KO) iPS cell lines. (A) Strategy for *SMAD3* deletion using CRISPR/Cas9. A schematic representation of the *SMAD3* gene is shown. Five sgRNAs (1–5) were designed upstream and five sgRNAs (6–10) downstream of *SMAD3* (indicated in green). The combination of one upstream and one downstream sgRNA could lead to a complete allele deletion. Primer binding sites for PCR and sequencing (pr3–pr6, indicated in purple) are indicated. (B) Surveyor assay to assess sgRNA efficiency. Lanes show a 100 bp marker and results of the Surveyor assay performed on PCR products (pr3–pr4 for sgRNAs 1–5, pr5–pr6 for sgRNAs 6– 10) from DNA of cells transfected with sgRNAs 1–10, with and without Surveyor nuclease treatment. The presence of cleavage products, smaller than the PCR product, indicates the formation of heteroduplexes due to CRISPR/Cas9-induced indels. sgRNA-4 and sgRNA-8 (indicated in red) were selected for further experiments. (C) PCR-based genotyping of the *SMAD3* KO cell line. The pr3–pr6 PCR only produces a product if *SMAD3* is deleted. The pr5–pr6 and pr3–pr4 PCRs generate a product only if the original *SMAD3* sequence (including possible indels) is present. The pr4–pr6 and pr3–pr5 PCRs yield a product only if the *SMAD3* allele is inverted. (D) Sanger sequencing to confirm *SMAD3* deletion and inversion in iPS cell line. Allele A and Allele B represent the two *SMAD3* alleles in the cell. The pr3–pr6 primers, located outside of *SMAD3*, only amplify a product if the entire *SMAD3* gene is deleted. The red line marks the Cas9 cut site.

### Transfections

HEK293 cells were seeded one day before transfection to reach ∼70% confluency. The next day, HEK293 cells were transfected with Lipofectamine 3000 according to the manufacturer’s protocol (ThermoFisher). All plasmids with sgRNAs were separately transfected into HEK293 cells to perform a Surveyor assay (see below).

Transfections to generate small insertions and deletions (indels) were performed with Lipofectamine 3000. The transfection mix with 3 ug of the eSpCas9(1.1)-T2A-EGFP with the selected sgRNA plasmid was added to a single-cell suspension of 2 million cells generated by Accutase treatment. Media was replaced after 24 hours. After two days cells were FACS sorted.

To generate *SMAD3* knock-out iPS cell lines, iPS cells were transfected with a combination of 2 plasmids; pSpCas9(BB)-2A-Puro (PX459) V2.0 with sgRNA (sgRNA4 and sgRNA8) using the Amaxa Human Stem Cell Nucleofector Kit 2 (Lonza).

### FACS + colony picking

iPS cells transfected with plasmid eSpCas9(1.1)-T2A-EGFP were dissociated with Accutase and run through a cell strainer two days after transfection. Cells were FACS sorted to be GFP positive and single cell. Next, cells were seeded at a density of 6,000 cells in a 6-well plate. After harvesting, cells were kept in mTeSR Plus with 10 mM Y-27672 ROCK inhibitor (Tocris, 1254). After a week colonies were picked under the microscope. iPS cells transfected with plasmid pSpCas9(BB)-2A-Puro (PX459) V2.0 were seeded at low density and colonies were picked under a microscope. The picked colonies were expanded and split into two wells on 24-well plates plates; one for DNA isolation and genotyping, and one to maintain cells.

### DNA isolation and PCR

Lysis buffer (100 mM Tris pH 8,5, 5mM EDTA pH 8,0, 200 mM NaCl, 0,2% SDS, 0,1 mg/ml proteinase K) was added to cells and it was incubated overnight at 37°C. Phenol:chloroform:isoamyl alcohol (25:24:1) was added in a 1:1 ratio to the lysed cells. After centrifugation, the aqueous phase was transferred to a new tube. DNA was then precipitated in ice-cold isopropanol and after centrifugation the pellet was dissolved in dH_2_O. DNA concentration was measured with Denovix DS-11 spectrophotometer. PCRs were performed with Pfu polymerase according to the manufacturer’s protocol (Promega, M7741) (primers: Supplementary Table 1), using 25 ng DNA, according to the following PCR protocol: 95°C for 2 min, 35 cycles of [95°C for 1 min, 60°C for 30 s, 72°C for 1 min], and a final extension at 72°C for 5 min.

### Surveyor assay

A Surveyor assay was performed on DNA isolated from HEK293 cells transfected with the individual plasmids to test the efficiency of sgRNAs introducing double strand breaks. The Surveyor Assay Mutation Detection Kit (IDT, 706020) was used according to the manufacturer’s protocol. For each target location one sgRNA was selected for iPS transfections.

### Genotyping

All cell lines were genotyped to confirm the introduction of the *SMAD3* variant. After DNA isolation and PCR, PCR products were purified with the QIAquick Gel Extraction Kit according to the manufacturer’s protocol (Qiagen, 28706). Purified products were submitted for Sanger sequencing to Macrogen Europe.

### Reprogramming of fibroblasts to iPS

iPS cell lines were reprogrammed by the Erasmus MC iPS core facility; Human dermal fibroblasts were transduced using CytoTune™-iPS 2.0 Sendai vectors at the appropriate multiplicity of infection (MOI) (ThermoFisher Scientific, A16517). Transduced fibroblasts were seeded on Matrigel coated plates and fed daily with mTeSR1 medium. Around day 21, undifferentiated colonies were picked. The clones were tested for the absence of CytoTune™ 2.0 Sendai reprogramming vectors at passages 7/8. To fully characterize the resulting iPS cells, all clones were assessed for the expression of undifferentiated stem cell markers by immunofluorescence and RT-PCR. Pluripotency was evaluated by differentiating the cells into the three embryonic germ layers: ectoderm, mesoderm and endoderm using the STEMdiff™ Trilineage Differentiation Kit (Stemcell Technologies, 05230) and examining marker expression. Both the original iPS cell lines and their differentiated lineages were analyzed by immunofluorescence and RT-qPCR to confirm the expression of germ-layer-specific markers. The official names of the generated cells are EMC178i, EMC179i, EMC173i. In this paper we refer to the cell lines as, iPS #1 p.R287W, iPS #2 p.R287W, and iPS #3 p.F248Pfs*62, respectively.

### iPS differentiation to SMC

iPS cells were differentiated into contractile SMCs as published [10]. iPS cells were seeded at a density of 200,000 single cells per well in a 6-well plate on day -2. Differentiation was started on day 0; see Figure 5A for media and supplements. The medium was replaced every 2 days. After differentiation, cells were cultured in DMEM with 5 ng/mL FGF and 5% serum in gelatin-coated dishes.

### Immunofluorescence

Cells were grown on coverslips and fixed with 4% paraformaldehyde in PBS for 15 minutes. After fixation, cells were permeabilized with PBS supplemented with 0.1% Triton-X-100 and blocked with PBS+ (PBS with 0.5% BSA and 0.15% glycine) for 30 minutes. Coverslips were incubated overnight at 4°C with primary antibodies (Supplementary Table 3) in PBS+. Coverslips were washed with PBS supplemented with 0.1% Triton-X-100 and briefly with PBS+ before incubation with Alexa Fluor secondary antibodies (1:1000, Molecular Probes) in PBS+ for one hour at room temperature. Coverslips were mounted in Vectashield with DAPI (H-1200, Vector Laboratories) and sealed with nail polish. Images were recorded with an Axio Imager D2 microscope (Zeiss).

### qPCR

RNA isolation was performed using the RNeasy Mini Kit (Qiagen). cDNA was synthesized using the iScript cDNA Synthesis Kit (Bio-Rad) according to the manufacturer’s protocol. For qPCR, 200 nM forward and reverse primers (Supplementary Table 4) and iQ™ SYBR^®^ Green Supermix (Bio-Rad) were combined with cDNA on the CFX384 system (Bio-Rad). The qPCR conditions were as follows: denaturation at 95°C for 3 min, followed by 40 cycles of denaturation at 95°C for 15 s, and annealing/extension at 60°C for 30 s. hGAPDH was used as the reference gene. Relative gene expression levels were determined using the comparative ΔΔCt method. ΔCT = CT(gene of interest) ™ CT(reference gene) and ΔΔCT = ΔCT(gene of interest) – average ΔCT(gene of interest, controls). The fold change is calculated as 2^-ΔΔCT.

### Western blotting

Cells were scraped in phosphate-buffered saline (PBS) supplemented with protease inhibitor cocktail (1:100, Roche, 11836145001) and phosphatase inhibitor cocktail (1:100, Sigma, P0044). Samples were lysed in equal volumes of 2x Laemmli buffer (4% SDS, 20% glycerol, 120mM Tris pH 6,8) supplemented with protease and phosphatase inhibitors. Lysates were passed through a 25G needle and heated to 65°C for 10 minutes. Protein concentrations were measured using the Lowry protein assay [11]. Equal amounts of protein were separated on an SDS-PAGE gel and transferred to an Immobilon-P polyvinylidene difluoride (PVDF) membrane (Millipore, IPVH00010). Membranes were blocked in PBS with 3% milk powder (Sigma, 70166) and 0.1% Tween-20 (Sigma, P1379). After blocking, the membranes were incubated overnight in blocking buffer with primary antibodies (Supplementary Table 5). Membranes were washed five times with PBS containing 0,1% Tween-20 and incubated for one hour at room temperature with horseradish peroxidase-conjugated secondary antibodies (1:2000, Jackson ImmunoResearch, 515-035-003 and 711-035-152). After washing, protein detection was performed using home-made enhanced chemiluminescence (ECL) substrate on an Amersham Imager 600 (GE Healthcare Life Sciences). Protein signal quantification was performed using Fiji software [12].

### Statistics

Data were corrected for outliers using the Grubbs’ test. Statistical analysis was performed with a non-parametric Mann-Whitney test. The standard deviation (SD) is shown in the figures. All analyses were performed using GraphPad Prism, version 8.

## Results

### Generation and validation of *SMAD3*-Indel iPS cell lines

To generate iPS cell lines with small insertions or deletions (indels) in the *SMAD3* gene, three different single guide RNAs (sgRNAs) (Figure 1A) were designed and cloned into the eSpCas9(1.1)-T2A-EGFP plasmid. Each plasmid was individually transfected in HEK293 cells, and the efficiency of double strand break (DSB) induction was assessed in a Surveyor assay. This assay detects sequence heterogeneity at the targeted locus by identifying mismatches that arise from imperfect repair following DSB formation. In a mixed population of transfected cells, PCR amplification of the target region generates a product containing both wild-type and indel-bearing sequences. Following denaturation and reannealing, heteroduplexes form and are subsequently cleaved by Surveyor nuclease, producing distinct fragment patterns on an agarose gel. All three sgRNAs successfully induced DSBs, as evidenced by the cleavage products, smaller than the 121 bp fragments that are expected when no DSBs are induced, observed on the gel (Figure 1B), with comparable efficiencies. sgRNA-A was selected for further experiments as it targeted a region near a known *SMAD3* P/LP variant within exon 6, a hotspot for pathogenic *SMAD3* variants in patients [13, 14, 1, 15, 16].

Plasmid eSpCas9(1.1)-T2A-EGFP carrying sgRNA-A was transfected into iPS cells, followed by fluorescence-activated cell sorting (FACS) to isolate eGFP-positive cells. These eGFP-positive cells were cultured, and individual colonies were expanded. Two clones (iPS #A indel and iPS #B indel) were selected, and the targeted genomic region was PCR-amplified using colony PCRs for the region around the sgRNA primers pr1 and pr2 (Figure 1C). Sanger sequencing of these PCR products confirmed the introduction of indels; a 4 bp deletion in one allele of iPS#A indel and a 34 bp deletion in one allele of iPS #B indel (Figure 1D).

### Generation and validation of iPS cells with a *SMAD3* gene deletion

To generate a knock-out of the *SMAD3* gene, ten different sgRNAs were designed; five targeting the upstream region and five targeting the downstream region of the *SMAD3* gene (Figure 2A). These sgRNAs were cloned into the pSpCas9(BB)-2A-Puro (PX459) V2.0 plasmid and transfected individually into control iPS cells to assess their efficiency in introducing double-strand breaks (DSBs). A Surveyor assay confirmed that most sgRNAs induced DSBs, as evidenced by cleaved PCR products, smaller than the PCR product without DSBs, visible on an agarose gel (Figure 2B).

Among the tested sgRNAs, sgRNA4 (targeting upstream of *SMAD3*) and sgRNA8 (targeting downstream) were selected for simultaneous transfection to induce a complete deletion of the *SMAD3* locus, as the Surveyor assay showed that they can properly introduce DSBs. Following transfection, individual colonies were isolated, and PCR screening identified one colony (iPS #C deletion) with a deleted *SMAD3* allele, as indicated by the presence of a PCR product using primers pr3-pr6. Additionally, the absence of amplification with primers pr5-pr6 and pr3-pr4 suggested that the original sequence was no longer intact (Figure 2C). Further PCR analysis revealed that a *SMAD3* allele was still present in the cell. PCR using multiple primer sets flanking the sgRNA target sites suggested that instead of being fully excised, one allele had undergone an inversion (Figure 2A and 2C). Sanger sequencing confirmed that iPS #C deletion contained one deleted allele and one inverted allele, with a single-base-pair deletion at the inversion junction (Figure 2D).

### Reprogramming of *SMAD3* patient fibroblasts into iPSCs and pluripotency validation

Human dermal fibroblasts of *SMAD3* patients were reprogrammed into iPS cells using Sendai virus-based delivery of reprogramming factors (Figure 3A). The resulting iPSC lines included iPS #1 (*SMAD3* p.R287W), iPS #2 (*SMAD3* p.R287W), and iPS #3 (*SMAD3* p.F248Pfs*62).

**Figure 3.**
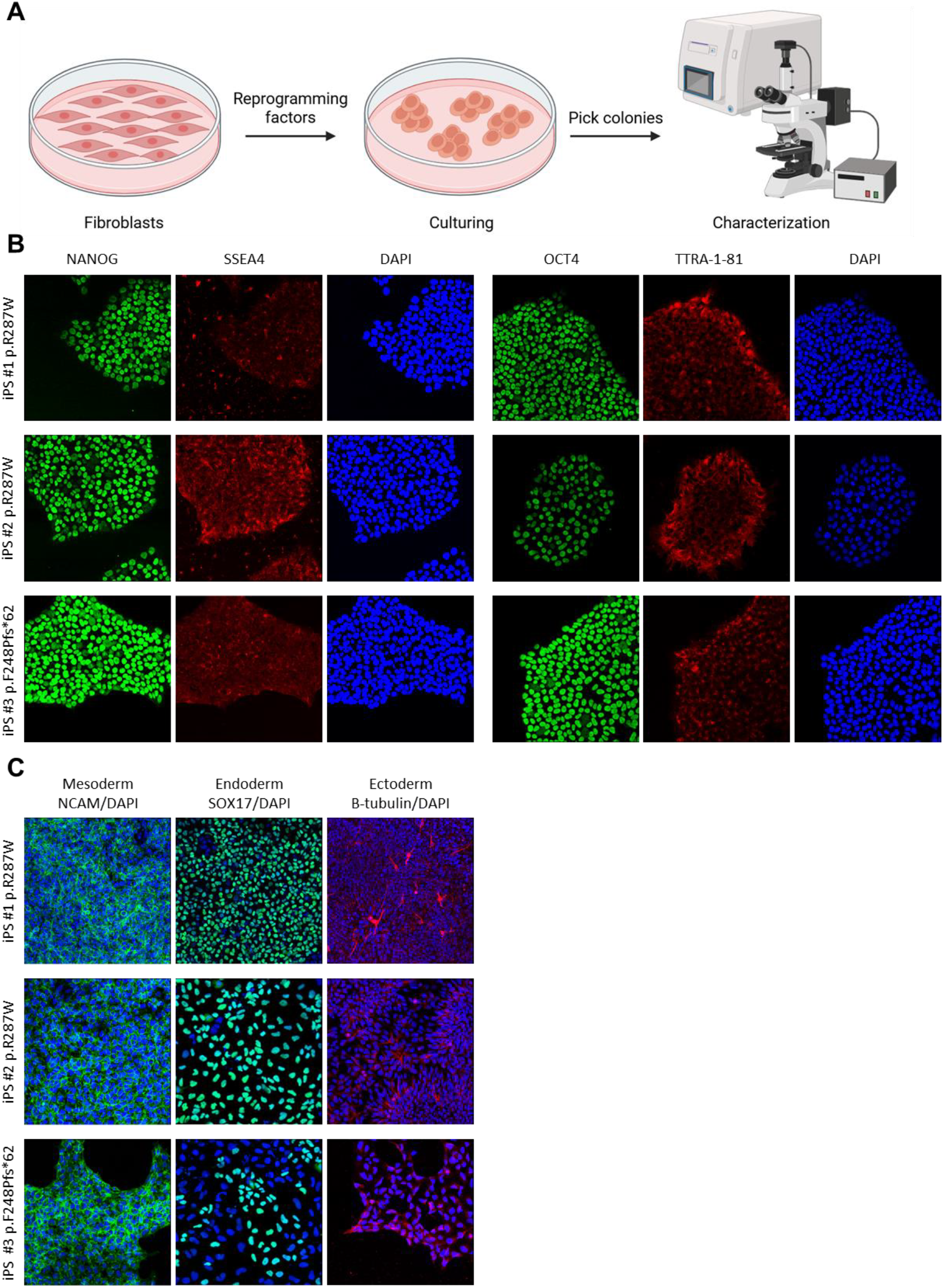
Generation and characterization of patient-derived iPS cell lines. (A) Reprogramming strategy from fibroblasts to iPS cells. Fibroblasts were transduced with Sendai virus carrying reprogramming factors. After several weeks of culture, individual colonies were picked and expanded. The resulting iPS cell lines were characterized to confirm successful reprogramming. (C) Immunofluorescent staining of pluripotency markers. Staining of patient-derived iPS cell lines for NANOG (green), SSEA4 (red), and DAPI (blue), as well as OCT4 (green), TRA-1-81 (red), and DAPI (blue) in iPS #1 (*SMAD3* p.R287W), iPS #2 (*SMAD3* p.R287W), and iPS #3 (*SMAD3* p.F248Pfs*62*). (C) Immunofluorescent staining of germ layer markers after differentiation. Patient-derived iPS cell lines (iPS #1 p.R287W, iPS #2 p.R287W, iPS #3 p.F248Pfs*62) were differentiated into mesoderm, endoderm, and ectoderm. Staining was performed for NCAM (mesoderm, green), SOX17 (endoderm, green), and β-tubulin (ectoderm, red). All images are overlays with DAPI staining (blue).

The *SMAD3* p.R287W variant, present in iPS #1 and iPS #2, is a point mutation that likely exerts a dominant-negative effect by disrupting protein complex formation [17]. In contrast, iPS #3 carries the *SMAD3* p.F248Pfs*62 variant, a frameshift variant known to trigger nonsense-mediated decay (NMD), leading to haploinsufficiency [2].

Pluripotency of the generated iPS cell lines was confirmed through immunofluorescence staining for the pluripotency markers NANOG, SSEA4, OCT4, and TRA-1-81 (Figure 3B), as well as quantitative PCR (qPCR) analysis for *OCT3/4, NANOG*, and *REX1* (Supplementary figure 1A-C). Furthermore, these iPSCs retained the capacity to differentiate into all three germ layers, as demonstrated by immunofluorescence staining for NCAM (mesoderm), SOX17 (endoderm), and β-tubulin (ectoderm) (Figure 3C), along with qPCR analysis of lineage-specific markers *Brachyury* (mesoderm), *FOXA2* (endoderm), and *PAX6* (ectoderm) (Supplementary figure 3D-F).

### SMAD3 protein expression in CRISPR-modified and patient-derived iPSCs

To assess SMAD3 protein expression, we performed Western blots on CRISPR/Cas9 generated iPS cell lines iPS #A indel, iPS #B indel, iPS #C deletion and patient derived iPS cell lines iPS #1 p.R287W, iPS #2 p.R287W, iPS #3 p.F248Pfs*62. SMAD3 protein was detected in all iPS cell lines (Figure 4A). In the CRISPR/Cas9-modified lines, expression appeared slightly reduced in iPS #A indel and iPS #B indel, though this reduction was not statistically significant (Figure 4A, B). In contrast, iPS #C deletion, which harbors one deleted *SMAD3* allele and one inverted allele, showed a significant decrease in SMAD3 protein levels (Figure 4A, B). In patient-derived iPS cell lines, SMAD3 expression was comparable to controls (Figure 4C, D).

**Figure 4.**
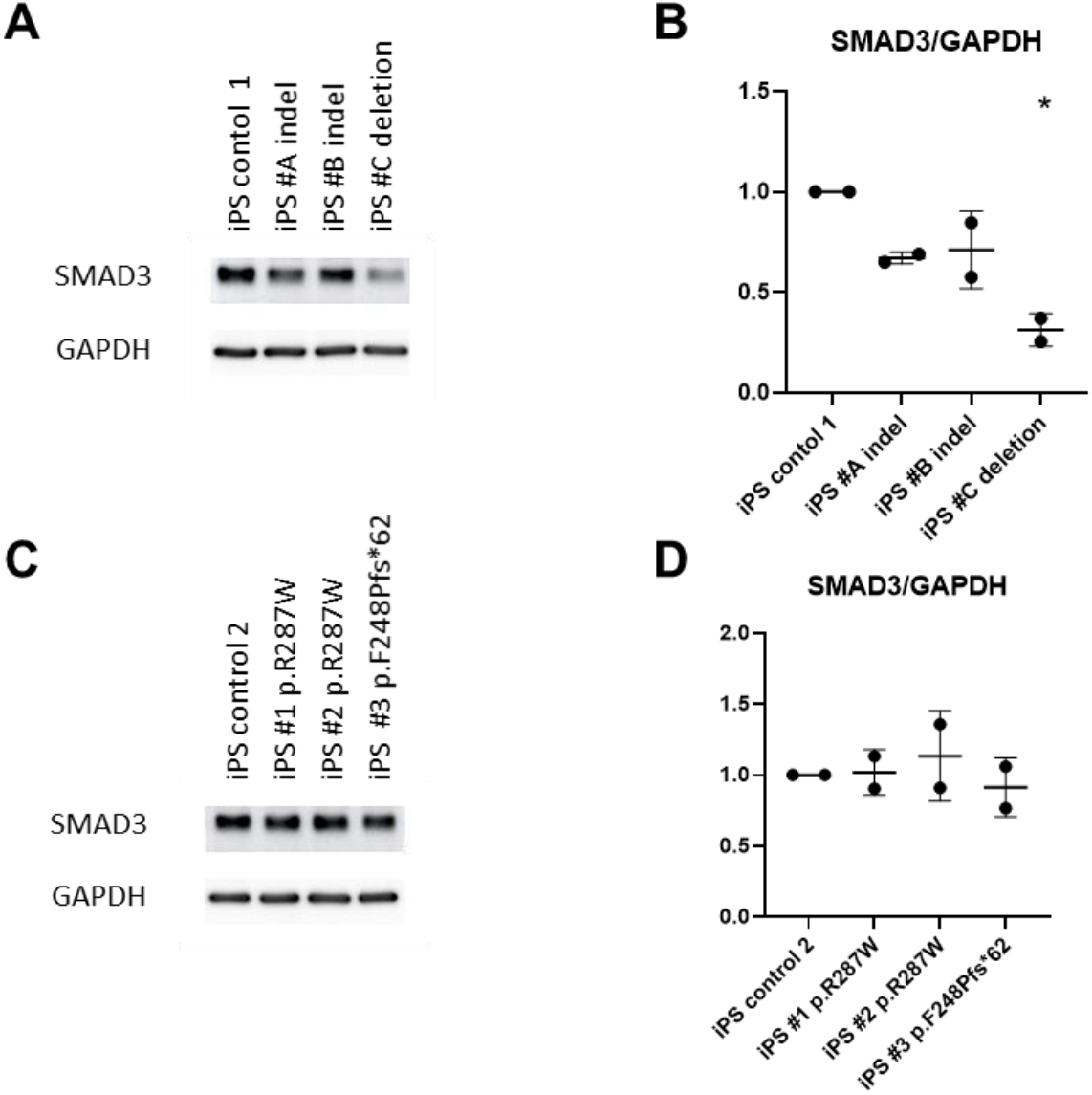
Expression of *SMAD3* in iPS cell lines. (A) Western blot analysis of *SMAD3* expression in CRISPR/Cas9-edited iPS cell lines. *SMAD3* protein levels were analyzed in iPS #A indel, iPS #B indel, and iPS #C deletion cell lines. GAPDH was used as a loading control. (B) Quantification of *SMAD3* protein levels in CRISPR/Cas9-edited iPS cells. Densitometric analysis of *SMAD3* expression from the western blot in (A). (C) Western blot analysis of *SMAD3* expression in patient-derived iPS cell lines. *SMAD3* protein levels were analyzed in iPS #1 (*SMAD3* p.R287W), iPS #2 (*SMAD3* p.R287W), and iPS #3 (*SMAD3* p.F248Pfs*62*). GAPDH was used as a loading control. (D) Quantification of *SMAD3* protein levels in patient-derived iPS cells. Densitometric analysis of *SMAD3* expression from the western blot in (C).

### Differentiation of iPSCs into smooth muscle cells for LDS3 modeling

Patient derived iPS (iPS #1 p.R287W, iPS #2 p.R287W, iPS #3 p.F248Pfs*62) and control iPS cells were differentiated into SMCs as this cell type is affected in LDS3, making it a relevant model for studying the molecular mechanism underlying the disease (Figure 5A). During differentiation, changes in morphology were visible with light microscopy (Figure 5B). After differentiation, we measured a loss of the iPS-specific Oct3/4, and a marked increase in the expression of SMC-specific markers, including SMA, MYH11, and SM22 as shown by Western blot analysis (Figure 5C). These results demonstrate that both control and *SMAD3*-mutant iPS cell lines are capable of differentiating into SMC-like cells.

**Figure 5.**
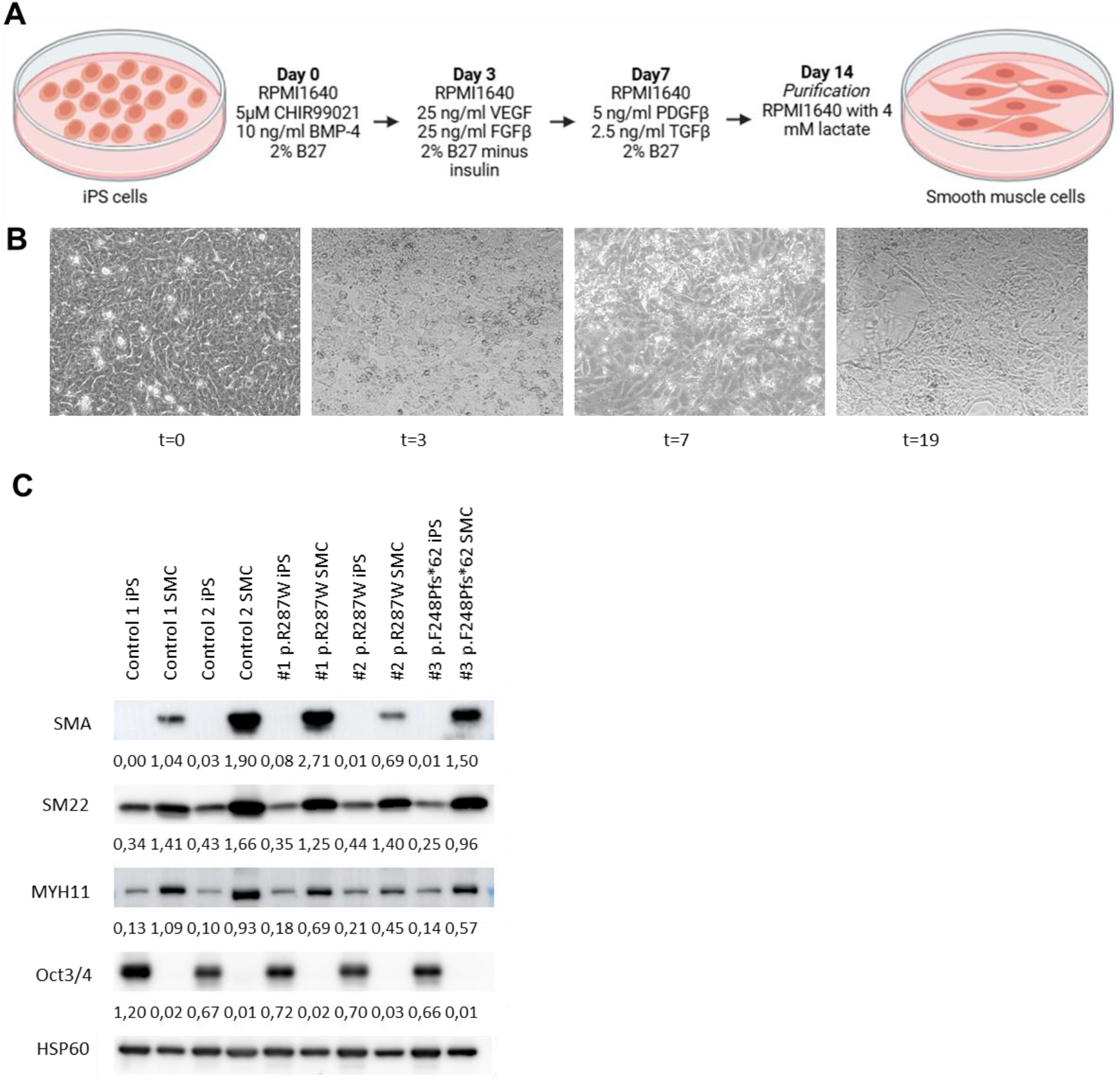
Differentiation of iPS cells into smooth muscle cells (SMCs). (A) Schematic representation of the differentiation protocol from iPS cells to SMCs. The sequential steps of differentiation are depicted. Culture medium was refreshed every two days. (B) Morphological changes during differentiation (control 1). Representative images of cells at days 0, 3, 7, and 19 of the differentiation process. (C) Western blot analysis of SMC and iPS markers. Protein expression of SMC markers (SMA, SM22, and MYH11) and the pluripotency marker OCT3/4 was analyzed in differentiated and undifferentiated iPS cell lines (iPS #1 *SMAD3* p.R287W, iPS #2 *SMAD3* p.R287W, iPS #3 *SMAD3* p.F248Pfs*62*). HSP60 was used as a loading control. Below each Western blot the quantification of protein levels is shown normalized to HSP60 levels.

## Discussion

IPS cells are a powerful model to study mechanisms of disease. In this study, we aimed to generate and validate iPS-based cell models to investigate Loeys-Dietz Syndrome type III (LDS3) caused by variants in the *SMAD3* gene. To achieve this, we employed two main approaches: CRISPR/Cas9-mediated gene editing in iPS cells and reprogramming of LDS3 patient fibroblasts carrying *SMAD3* P/LP variants.

Multiple *SMAD3* iPS cell lines were successfully generated. The mutations introduced (4 and 34 bp indels) likely lead to nonsense-mediated decay (NMD), as the resulting sequence runs out of frame, producing an early stop codon. This was consistent with the observed reduction in SMAD3 protein expression in these iPS cell lines

Although our goal was to completely knock out *SMAD3* by deleting both alleles, we instead created a cell line with one *SMAD3* allele fully deleted and the other allele inverted. We observed a partially reduced SMAD3 protein expression, which aligns with the genetic nature of the disease, where most *SMAD3* P/LP variants are heterozygous. Thus, the model with a significant but not complete reduction of SMAD3 may better mimic the disease phenotype. Importantly, because this cell line was generated by CRISPR/Cas9, the original cell line is an isogenic control, allowing us to study the effect of the specific genetic changes without interference from other genetic or environmental factors. This makes it a very useful model for differentiation into SMC and conducting functional cellular assays, such as assessing SMC markers, and extracellular (ECM) formation, both of which are affected in patient-derived *SMAD3* VSMCs and myofibroblasts [2, 17]. We hypothesize that the allele was cleaved, inverted, and then integrated into the genome through non-homologous end joining (NHEJ) repair. At one junction, this repair resulted in a 1 bp deletion. This explanation is consistent with findings from previous studies using two sgRNAs [18].

We also validated iPS cell lines reprogrammed from *SMAD3* patient fibroblasts. These lines expressed pluripotency markers (*NANOG, OCT4, SSEA4, TTRA-1-81*) and successfully differentiated into ectoderm, mesoderm, and endoderm lineages. Although SMAD3 has been shown to co-occupy OCT4 targets during reprogramming and to contribute to reprogramming efficiency, it is not essential for the process [19], which is also confirmed by our own data. Notably, all *SMAD3* patient iPS cell lines used for reprogramming contained at least one functional *SMAD3* allele, which likely explains the successful differentiation into SMCs.

Western blot analysis showed that SMAD3 protein expression was not significantly reduced in the patient-derived iPS lines. In iPS #1 p.R287W and iPS #2 p.R287W, the mutated *SMAD3* allele is still transcribed and translated, though the resulting protein is likely non-functional. As the antibody for SMAD3 recognizes a part of SMAD3 where this P/LP variant is not located, it is likely that the antibody recognizes healthy and mutated SMAD3 equally well, which can explain the detection of normal SMAD3 levels. Similar results were shown in fibroblasts [2, 17]. However, the P/LP variant in iPS #3 p.F248Pfs*62 was shown to result in NMD and the fibroblasts that were used to generate this iPS cell line were shown to have a reduced SMAD3 protein expression [2, 17]. An explanation for this could be the fact that SMAD3 expression is specific to differentiated cell lines such as VSMCs and fibroblasts, and therefore, SMAD3 protein expression might be lower in iPS cells. As there is still one healthy *SMAD3* allele present, the expression of this allele alone could suffice to account for SMAD3 protein levels in iPS cells. It could also be that during reprogramming only cells with a relatively high SMAD3 protein expression survive. Another option is that several passages are needed after differentiation before the SMCs are fully differentiated and therefore to validate SMAD3 protein expression properly.

Differentiation of *SMAD3* patient-derived iPS cells into SMCs is crucial for studying the aortic phenotype associated with LDS3. Previous studies in VSMCs and myofibroblasts have shown molecular and cellular changes, such as altered expression of VSMC markers and ECM formation, in response to *SMAD3* variants. In this study, we successfully differentiated the *SMAD3* iPS cells into SMCs, validated by the expression of SMC-specific markers. Moving forward, functional assays in these models can explore transcriptional activation of TGF-β signaling, ECM formation, and VSMC marker expression which are expected to be impacted in LDS3 [20, 17]. In this study we focused on the generation of different *SMAD3* iPS-based cell models and differentiated them to SMCs, validated by expression of SMC-specific protein markers. While this study focused on differentiating iPS cells into SMCs, it is important to note that LDS3 also frequently involves joint issues, such as osteoarthritis. iPS cell models not only allows to study *SMAD3* variants in SMCs, but also provide the opportunity to explore the impact of these mutations on other relevant cell types, such as chondrocytes to study osteoarthritis. Future research could expand the model to include iPS-derived chondrocytes with *SMAD3* variants, which would provide further insight into the broader effects of *SMAD3* variants on connective tissue in LDS3.

In conclusion, this study demonstrates the potential of iPS-based models for studying SMAD3 variants. iPS cells offer a versatile platform for disease modeling due to their ease of generation, potential for isogenic controls, and ability to differentiate into multiple cell types. By focusing on the differentiation of iPS cells into SMCs, we have created a relevant model for studying LDS3, which can be further expanded to explore additional cell types and functional assays.

## Supporting information

Suppplementary

## Acknowledgements

We would like to thank the department of Clinical Genetics, laboratory diagnostics, for the preparation and storage of patient fibroblasts.

