## Supplementary material for "Generation and characterization of Induced Pluripotent Stem Cell models with *SMAD3* mutations to study the molecular mechanism of Loeys-Dietz Syndrome Type III aneurysm formation": Suppplementary

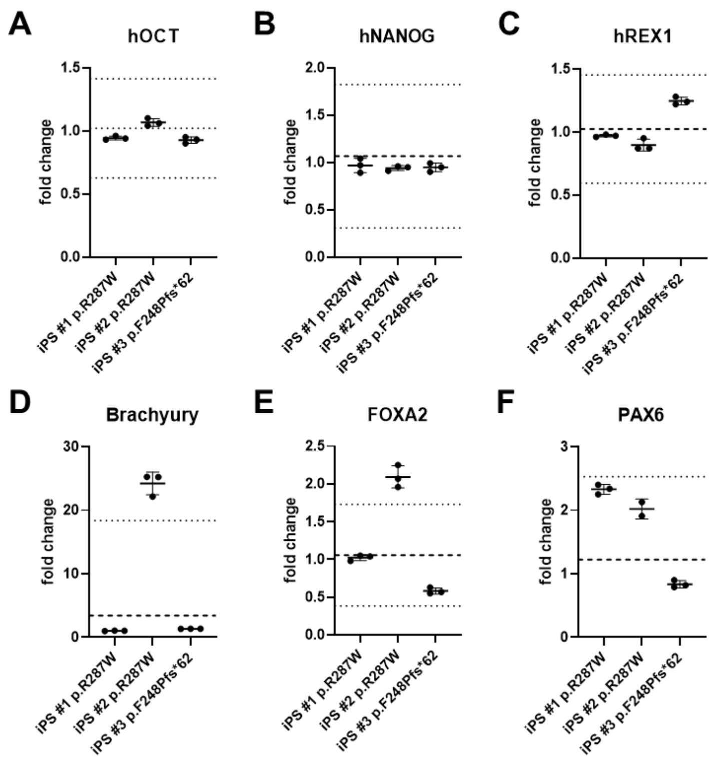


**Supplementary figure 1. qPCRs generation iPS patient cell lines**
(A-C) Relative mRNA expression of pluripotency markers in patient-derived iPS cell lines. Quantitative PCR (qPCR) analysis of pluripotency-associated genes in iPS #1 (*SMAD3* p.R287W), iPS #2 (*SMAD3* p.R287W), and iPS #3 (*SMAD3* p.F248Pfs*62*). Expression levels are shown for: (A) *hOCT4*, (B) *hNANOG*, (C) *hREX1*. (D-F) Relative mRNA expression of germ layer-specific markers in differentiated patient-derived iPS cells. qPCR analysis of lineage-specific genes in iPS #1 (*SMAD3* p.R287W), iPS #2 (*SMAD3* p.R287W), and iPS #3 (*SMAD3* p.F248Pfs*62*) after differentiation into the three germ layers. Expression levels are shown for: (D) *Brachyury* (mesoderm), (E) *FOXA2* (endoderm), (F) *PAX6* (ectoderm).

**Supplementary table 1: sgRNA sequences**

| Location | Name sgRNA | Sequence sgRNA | |
|---|---|---|---|
| Exon 6 | sgRNA A | 5’-caccgATGCAGCAGTGGAGCTGACA-3’ | 5’-aaacTGTCAGCTCCACTGCTGCATC-3’ |
| | sgRNA B | 5’-caccGGAGACACATCGGTATGGGG-3’ | 5’-aaacCCCCATACCGATGTGTCTCC-3’ |
| | sgRNA C | 5’-caccGACACGGAGACACATCGGTA-3’ | 5’-aaacTACCGATGTGTCTCCGTGTC-3’ |
| Upstream of SMAD3 gene | sgRNA1 | 5’-caccGGGCTTAGACACCGGCTAGT-3’ | 5’-aaacACTAGCCGGTGTCTAAGCCC-3’ |
| | sgRNA2 | 5’-caccgTAGCTCCTGAAAACCGCCGA-3’ | 5’-aaacTCGGCGGTTTTCAGGAGCTAc-3’ |
| | sgRNA3 | 5’-caccgTGGATTCCGCCACACCTCAC-3’ | 5’-aaacGTGAGGTGTGGCGGAATCCAc-3’ |
| | sgRNA4 | 5’-caccgATTCAGGGGCGTCTGCCCGC-3’ | 5’-aaacGCGGGCAGACGCCCCTGAATc-3’ |
| | sgRNA5 | 5’-caccgTCCTAAATACCCGCCTGAAA-3’ | 5’-aaacTTTCAGGCGGGTATTTAGGAc-3’ |
| Downstream of SMAD3 gene | sgRNA6 | 5’-caccgCTAGGTAGGAAGAGCCCGCA-3’ | 5’-aaacTGCGGGCTCTTCCTACCTAGc-3’ |
| | sgRNA7 | 5’-caccgCTACCTAGAACGCCCTTCCA-3’ | 5’-aaacTGGAAGGGCGTTCTAGGTAGc-3’ |
| | sgRNA8 | 5’-caccgCTTGTGAGCTGGATAGACTT-3’ | 5’-aaacAAGTCTCTCCAGCTCACAAGc-3’ |
| | sgRNA9 | 5’-caccgCCCTCTCAGAACATACTGAT-3’ | 5’-aaacATCAGTATGTTCTGAGAGGGc-3’ |
| | sgRNA10 | 5’-caccgCCTAGACAGAGGGGTGTTTC-3’ | 5’-aaacGAAACACCCCTCTGTCTAGGc-3’ |

**Supplementary table 2: Primer sequences used to analyze and validate CRISPR/Cas9 generated iPS cell lines**

| Location | Primer | Forward/ reverse | Primer sequence |
|---|---|---|---|
| Upstream of exon 6 SMAD3, before sgRNAs | #1 | Forward | 5’-ATGACTGTGGATGGCTTCACC-3’ |
| Downstream of exon 6 SMAD3, after sgRNAs | #2 | Reverse | 5’-TAAGGATGGACGCAAGGTTCC-3’ |
| Upstream of SMAD3, before sgRNAs | #3 | Forward | 5’-AGACTGAATCAAAGGGCAGCA-3’ |
| Upstream of SMAD3, after sgRNAs | #4 | Reverse | 5’-AACAGGGTGTTTGTTGCTGG-3’ |
| Downstream of SMAD3, before sgRNAs | #5 | Forward | 5’-CCGCTGTTCCAGTGTGTCTT-3’ |
| Downstream of SMAD3, after sgRNAs | #6 | Reverse | 5’-AGATGACTGGAGACGCACC-3’ |

**Supplementary table 3: Primary antibodies used for immunofluorescence**

| Primary antibody | Dilution | ES/mesoderm/ectoderm/endoderm | Manufacturer | Catalog number |
|---|---|---|---|---|
| SSEA4 | 1:75 | ES | Invitrogen | 41-4000 |
| NANOG | 1:75 | ES | Abcam | Ab21624 |
| TRA-1-81 | 1:250 | ES | Abcam | Ab16289 |
| OCT4 | 1:300 | ES | Abcam | Ab19857 |
| NCAM | 1:100 | Mesoderm | R&D Systems | AF2408 |
| SOX17 | 1:50 | Endoderm | BioTechne | AF1924-SP |
| B-tubulin | 1:1000 | Ectoderm | Sigma | T8660 |

**Supplementary table 4: Primer sequences used for qPCR**

| Gene | ES/mesoderm /ectoderm/ endoderm/ intern control | Forward primer | Reverse primer |
|---|---|---|---|
| hOCT3/4 | ES | 5'-GACAGGGGGAGGGGAGGAGCTAGG-3' | 5'-CTTCCCTCCAACCAGTTGCCCCAAAC-3' |
| hNANOG | ES | 5'-CAGCCCTGATTCTTCCACCAGTCCC-3' | 5'-CGGAAGATTCCCAGTCGGGTTCACC-3' |
| hREX1 | ES | 5'-CAGATCCTAAACAGCTCGCAGAAT-3' | 5'-GCGTACGCAAATTAAAGTCCAGA-3' |
| hBrachyury | Mesoderm | 5'-GGATGAAGGCTCCCGTCTC-3' | 5'-GCTGTGATCTCCTCGTTCTGATA-3' |
| hFOXA2 | Endoderm | 5'-TACAGGCGCAGCTACACGCACGCAAAG-3' | 5'-GCGGGGCACCTTCAGGAAACAGTCGT-3' |
| hPAX6 | Ectoderm | 5'-TTTGCCCGAGAAAGACTAGC-3' | 5'-CATTTGGCCCTTCGATTAGA-3' |
| hGAPDH | Intern control | 5'-GCCAAAAGGGTCATCATCTC-3' | 5'-GGTGGTGCAGGAGGCATT-3' |

**Supplementary table 5: Primary antibodies used for Western blotting**

| Primary antibody | Predicted kDA | Dilution | Synthetic/contractile/iPS / loading control | Manufacturer | Catalog number |
|---|---|---|---|---|---|
| Rabbit α-SMAD3 IgG | 48 | 1:1000 | - | Abcam | Ab28379 |
| Mouse α-GAPDH IgG1 | 40 | 1:5000 | Loading control | Abcam | Ab8245 |
| Rabbit α-SM22 IgG | 23 | 1:2000 | Contractile | Abcam | Ab14106 |
| Mouse α-SMA IgG2a | 42 | 1:10.000 | Contractile | Abcam | Ab7817 |
| Rabbit α-MYH11 IgG | 227 | 1:1000 | Contractile | Abcam | Ab53219 |
| Goat α-Oct-3/4 IgG | 46 | 1:1000 | iPS | Santa Cruz Biotechnology | sc-8628 |
| Rabbit α-HSP60 IgG | 61 | 1:10.000 | Loading control | GeneTex | GTX110089 |
